# Phylogenomic analysis of natural products biosynthetic gene clusters allows discovery of arseno-organic metabolites in model streptomycetes

**DOI:** 10.1101/020503

**Authors:** Pablo Cruz-Morales, Johannes Florian Kopp, Christian Martinez-Guerrero, Luis Alfonso Yáñez-Guerra, Nelly Selem Mojica, Hilda Ramos-Aboites, Jörg Feldmann, Francisco Barona-Gómez

## Abstract

Natural products from microbes have provided humans with beneficial antibiotics for millennia. However, a decline in the pace of antibiotic discovery exerts pressure on human health as antibiotic resistance spreads, a challenge that may better faced by unveiling chemical diversity produced by microbes. Current microbial genome mining approaches have revitalized research into antibiotics, but the empirical nature of these methods limits the chemical space that is explored.

Here, we address the problem of finding novel pathways by incorporating evolutionary principles into genome mining. We recapitulated the evolutionary history of twenty-three enzyme families previously uninvestigated in the context of natural product biosynthesis in *Actinobacteria,* the most proficient producers of natural products. Our genome evolutionary analyses where based on the assumption that expanded-repurposed enzyme families-from central metabolism, occur frequently and thus have the potential to catalyze new conversions in the context of natural products biosynthesis. Our analyses led to the discovery of biosynthetic gene clusters coding for hidden chemical diversity, as validated by comparing our predictions with those from state-of-the-art genome mining tools; as well as experimentally demonstrating the existence of a biosynthetic pathway for arseno-organic metabolites in *Streptomyces coelicolor* and *Streptomyces lividans*, using gene knockout and metabolite profile combined strategy. As our approach does not rely solely on sequence similarity searches of previously identified biosynthetic enzymes, these results establish the basis for the development of an evolutionary-driven genome mining tool that complements current platforms. We anticipate that by doing so real ‘chemical dark matter’ will be unveiled.

## Introduction

Natural products (NPs) are a diverse group of specialized metabolites with adaptive functions, which include antibiotics, metal chelators, enzyme inhibitors, signaling molecules, amongst others (Traxler and Kolter, 2015). In bacteria, their biosynthesis is directed by groups of genes, referred to as Biosynthetic Gene Clusters (BGCs), usually encoded within a single locus, which allows for the concerted expression of biosynthetic enzymes, regulators, transporters and resistance genes (Diminic et al, 2014). Biosynthetic gene cluster formation has actually eased cloning of complete pathways and proposing experimentally verifiable *in silico* predictions of BGCs. Indeed, a fundamental principle of current bioinformatics approaches used for the discovery of novel NPs, commonly referred to as genome mining of NPs, rely in the assumption that once an enzyme is unequivocally linked to the production of a given metabolite, genes in the surroundings of its coding sequence are associated with its biosynthesis (Medema and Fischbach, 2015). This functional annotation approach, from genes to metabolites, has led to comprehensive catalogs of putative BGCs directing the synthesis of an ever-growing universe of metabolites (Medema et al, 2015; Hadjithomas et al, 2015).

NPs are also a rich source of compounds that have found pharmacological applications, as highlighted by the latest Nobel Prize in Physiology or Medicine awarded to researchers due to their contributions surrounding the discovery and use of NPs to treat infectious diseases. Indeed, in the context of increased antibiotic resistance, genome mining has revitalized the investigation into NP biosynthesis and their mechanisms of action (Demain, 2014; Harvey et al, 2015). In contrast with pioneering studies based on activity-guided screenings of NPs, current efforts based on genomics approaches promise to turn the discovery of NP drugs into a chance-free endeavor (Schreiber, 2005; Bachmann et al, 2014; Demain, 2014). Evidence supporting this possibility has steadily increased since ECO4601, a farnesylated benzodiazepinone discovered using genome mining approaches, which entered into human clinical trials more than a decade ago (Gourdeau et al, 2007).

Early genome mining approaches built up from the merger between a wealth of genome sequences and an accumulated biosynthetic empirical knowledge, mainly surrounding Polyketide Synthases (PKS) and Non-Ribosomal Peptide Synthetases (NRPSs) (Conway and Boddy, 2013; Ichikawa et al, 2013). These approaches can be classified as: (i) chemically-driven, where the discovery of the biosynthetic gene cluster is elucidated based on a fully chemically characterized ‘orphan’ metabolite (Barona-Gómez et al, 2004); or (ii) genetically-driven, where known sequences of protein domains (Lautru et al, 2005) or active-site motifs (Udwary et al, 2007) help to identify putative BGCs and their products. The latter relates to the term ‘cryptic’ BGC, defined as a locus that has been predicted to direct the synthesis of a NP, but which remains to be experimentally confirmed (Challis, 2008).

Decreasing costs of sequencing technologies has dramatically increased the number of putative BGCs. In this context, genome mining of NPs can help to prioritize strains on which to focus for further investigation (Rudolf et al, 2015; Shen et al, 2015). During this process, based on a priori biosynthetic insights, educated guesses surrounding PKS and NRPS can be put forward. In turn, such efforts increase the likelihood of discovering interesting chemical and biosynthetic variations. Moreover, biosynthetic logics for a growing number of NP classes, such as phosphonates (Metcalf and van der Donk, 2009; Ju et al, 2013), are complementing early NRPS/PKS-centric approaches. In contrast, finding novel chemical scaffolds, expected to be synthesized by cryptic BGCs, remains a challenging task. Therefore, with the outstanding exception of ClusterFinder (Cimmermancic et al, 2014), which uses Pfam domain pattern-based predictions, most genome mining methods are focused in known classes of NPs, hampering our ability to discover chemical novelty (Medema & Fischbach, 2015).

In this work we address the problem of finding novel pathways by genome mining, by means of integrating three evolutionary concepts related to emergence of NP biosynthesis. First, we assume that new enzymatic functions evolve by retaining their reaction mechanisms, while expanding their substrate specificities (Gerlt and Babbitt, 2001). In consequence, this process expands enzyme families. Second, evolution of contemporary metabolic pathways frequently occurs through recruitment of existing enzyme families to perform new metabolic functions (Caetano-Anollés et al, 2009). In the context of NP biosynthesis, the canonical example for this would be fatty acid synthetases as the ancestor of PKSs (Jenke-Kodama et al, 2005). Consequently, the correspondence of enzymes to either central or specialized metabolism, typically solved through detailed experimental analyses, could also be determined through phylogenomics. Third, BGCs are rapidly evolving metabolic systems, consisting of smaller biochemical sub-systems or ’sub-clusters’, which may have their origin in central metabolism (Vining, 1992; Firn and Jones, 2009 Medema et al, 2015).

These three evolutionary principles were formalized as a functional phylogenomics approach (Figure 1), which has been previously referred to as EvoMining (Medema & Fischbach, 2015). This approach leads to the identification of expanded, repurposed enzyme families, with the potential to catalyze new conversions in specialized metabolism. Using this approach we predicted several new potential pathways including the first ever reported family of BGCs for arseno-organic metabolites. Experimental evidence for arseno-organic metabolites produced by model actinomycetes *Streptomyces coelicolor* A3(2) and *Streptomyces lividans* is provided. As our approach does not rely solely on sequence similarity searches of previously identified NP biosynthetic enzymes, these results establish the basis for the development of an evolutionary-driven genome mining tool that complements current platforms.

**Figure 1.**
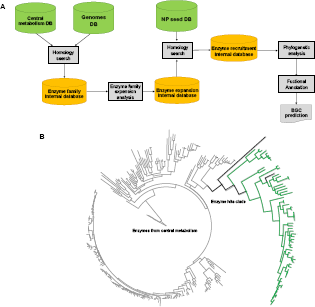
Phylogenomic analysis for the recapitulation of the evolution of NP biosynthesis. **A.** Bioinformatic workflow: the three input databases, as discussed in the text, are shown in green. Internal databases are shown in yellow, whereas grey boxes depict processes. **B.** An example of a phylogenetic tree for the recapitulation of the evolution of NP biosynthesis (**Tree S4**) using the case of 3-phosphoshikimate-1-carboxyvinyl transferase family. Gray branches include homologs related to central metabolism and their topology resembles that of a species guide tree (**Tree S1**). The Enzyme hits clade is highlighted: branches in black mark homologs from the Seed NP DB, and green branches indicate new enzyme hits. A detailed annotation of the clade is shown in figure 2C.

## Materials and methods

### Database integration

Genome DB: Complete and draft genomes of 230 members of the *Actinobacteria phylum* (supplementary table 1) were retrieved from GenBank either as single contigs or as groups of contigs in GenBank format, amino acid and DNA sequences were extracted from these files using in-house made scripts. Central metabolism DB: the amino acid sequences from the proteins involved in central metabolism were obtained from a database that we assembled for a previous enzyme expansion assessment (Barona-Gómez et al, 2012). The databases included a total of 339 queries for nine pathways, including amino acid biosynthesis, glycolysis, pentose phosphate pathway and tricarboxylic acids cycle (supplementary table 2) these sequences were obtained from the genome scale metabolic reconstructions of three actinobacterial model strains *S. coelicolor* (Borodina et al, 2005) *Mycobacterium tuberculosis* (Jamshidi and Palsson, 2007) and *Corynebacterium glutamicum* (Kjeldsen and Nielsen, 2009). Seed NP database: NRPS and PKS BGCs were obtained from the DoBISCUIT and ClusterMine 360 databases (Conway and Boddy, 2013; Ichikawa et al, 2013). BGCs for other NP classes were collected from available literature (supplementary table 3). Annotated GenBank formatted files were downloaded from the GenBank database to assemble a database that included 226 BGCs.

### Phylogenomics analysis pipeline

The sequences Central Metabolism DB were used as queries to retrieve members of each enzyme family from the Genomes DB using BlastP (Altschul et al 1990), with an e-value cutoff of 0.0001 and a bit core cutoff of 100. The average number of homologs of each enzyme family per genome and the standard deviation were calculated to establish a cutoff to identify and highlight significant expansion events (supplementary table 4). An enzyme family expansion was scored if the number of homologs in at least one genome was higher than the average number of homologs identified from each BLAST search. The next step was to BLAST sequences in the expanded families using amino acid sequences from the Seed NP DB as queries using an e-value cutoff of 0.0001 and a bit score cutoff of 100. When matches were identified, these represented candidate recruitment events.

The homologs found in known BGCs were added as seeds to the sequences from the expanded enzyme families with recruitments for future clade identification and labeling. The bidirectional best hits with the sequences in the Central Metabolism DB were identified and tagged using in-house scripts to distinguish central metabolic orthologs from other homologs that resulted from expansion events. These sets of sequences were aligned using Muscle version 3.8.31 (Edgar, 2004). The alignments were inspected and curated manually using JalView 2.8 (Waterhouse et al, 2009). The curated alignments were used for phylogenetic reconstructions, which were estimated using MrBayes 3.2.3 (Ronquist et al, 2012) with the following parameters: aamodelpr=mixed, samplefreq=100, burninfrac=0.25 in four chains and for 1000000 generations. The identification of new recruitments (hits) was done by visual inspection of each phylogentic tree. The homologs resulting from expansion events located in clades where at least one homolog from the Seed NP DB was found were selected and the regions of approximately 80 Kbs flanking their coding sequences were retrieved from GenBank formatted files using a perl Script. These contigs were annotated using the stand alone version of antiSMASH 3.0 (Weber et al, 2015) with the following options: -inclusive –cf_threshold 0.7 and RAST (Aziz et al, 2008). Further analysis of annotated contigs was done manually using the Artemis Genome Browser (Carver et al, 2012).

### Mutagenesis analysis

*S. coelicolor* (SCO6819) *and S. lividans* 66 (SLI_1096) knock-out mutants were constructed using in-frame PCR-targeted gene replacement of their coding sequences with an apramycin resistance cassette (*acc(3)*IV) (Gust et al, 2003). The plasmid pIJ773 was used as template to obtain a mutagenic cassette containing the apramycin resistance marker by PCR amplification with the primers reported in supplementary table 9. The mutagenic cassettes were used to disrupt the coding sequences of the genes of interest from the cosmid clone 1A2 that spans from SCO6971 to SCO6824 (Redenbach et al, 1996). Given the high sequence identity between the regions covered by cosmid 1A2 with the orthologous region in *S. lividans*, this cosmid clone was also used for disruption of SLI_1096. The gene disruptions were performed using the Redirect system (Gust et al, 2003). Double cross-over ex-conjugants were selected using apramycin resistance and kanamycin sensitivity as phenotypic markers. The genotype of the clones was confirmed by PCR. The strains and plasmids of the Redirect system were obtained from the John Innes Centre (Norwich, UK).

### RT-PCR analysis

The *S. lividans* 66 wild type strain was grown on 0 and 3; 0 and 300; 500 and 3; 500 and 300 µM of Arsenate and KH_2_PO_4_ respectively in solid modified R5 media for eight days. Mycelium collected from plates was used for RNA extraction with a NucleoSpin RNA II kit (Macherey-Nagel). The RNA samples were used as template for RT-PCR using the one step RT-PCR kit (Qiagen) (2ng RNA template for each 40 µl reaction). The housekeeping sigma factor *hrdB* (SLI_6088) was used as a control.

### LC-MS metabolite profile analysis

The SLI_1096 and SCO6818 minus mutants were grown on modified R5 medium (K_2_SO_4_ 0.25 g; MgCl_2_-6H_2_0 10.12 g; glucose 10 g; casamino acids 0.1 g; TES buffer 5.73 g; trace element solution (Kieser et al, 2000) 2 mL; agar 20 g) supplemented with a gradient of KH_2_PO_4_ and Na_3_AsO_4_ ranging from 3 to 300 µM and 0 to 500 µM, respectively. Induction of the arseno-organic BGC in both strains was detected in the condition were phosphate is limited and arsenic is available. Therefore, modified R5 liquid media supplemented with 3µM KH_2_PO_4_ and 500 µM Na_3_AsO_4_ was used for production of arseno-organic metabolites, and the cultures were incubated for 14 days in shaken flasks with metal springs for mycelium dispersion at 30° C. The mycelium was obtained by filtration, and the filtered mycelium was washed thoroughly with deionized water and freeze-dried. The samples were extracted overnight twice with MeOH/DCM (1:2). The extracts were combined and evaporated to dryness, and the dry residues were re-dissolved in 1 mL of MeOH (HPLC-Grade) and injected to the HPLC. The detection of organic arsenic species was achieved by online-splitting of the HPLC-eluent with 75% going to ESI-Orbitrap MS (Thermo Orbitrap Discovery) for accurate mass analysis and 25% to ICP-QQQ-MS (Agilent 8800) for the detection of arsenic. For HPLC, an Agilent Eclipse XDB-C18 reversed phase column was used with a H_2_O /MeOH gradient (0-20 min: 0-100% MeOH; 20-45 min: 100% MeOH; 45-50 min: 100% H_2_O). The ICP was set to oxygen mode and the transition ^75^As^+^ > (^75^As^16^O)^+^ (Q1: *m/z* = 75, Q2: *m/z* = 91) was observed. The correction for carbon enhancement from the gradient was achieved using a mathematical approach as described previously (Amayo et al, 2011). The ESI-Orbitrap-MS was set to positive ion mode in a scan range from 250-1100 amu. Also, MS2-spectra for the major occurring ions were generated.

8

## Results and discussion

### Evolutionary recapitulation of expanded and repurposed enzyme families

The proposed model for the evolution of NPs BGCs, based in the relationships between central metabolic enzymes and NP BGCs, was investigated through systematic and comprehensive phylogenomics (Figure 1A). The inputs for this analysis were formalized as three databases: (i) a genomes database consisting of 230 genome sequences belonging to the phylum *Actinobacteria,* which includes the most proficient producers of NPs, e.g. the genus *Streptomyces* (Genomes DB; Supplementary table 1); (ii) the amino acid sequences of central metabolic enzymes, belonging to nine selected pathways known to provide precursors for the synthesis of NPs (Central Metabolism DB, Supplementary table 2). These pathways, which have been shown to have enzymes that suffer from expansions with taxonomic resolution in *Actinobacteria* (Barona-Gómez et al, 2012), include glycolysis, tricarboxylic acid cycle, pentose phosphate pathway and the biosynthetic pathways for the proteinogenic amino acids. In total, these pathways consist of 106 enzyme families that were retrieved from well curated and experimentally validated genome-scale metabolic model reconstructions of model *Actinobacteria*, e.g. *Streptomyces coelicolor*; and (iii) a NP-related enzymes database including amino acid sequences from 226 known actinobacterial BGCs manually extracted from the literature, which provided a proof of concept limited universe (Seed NP DB, Supplementary table 3).

For each enzyme family, enzyme expansions were identified when the number of orthologs detected for a query sequence was greater than the average number detected in all genomes plus one standard deviation (Supplementary table 4). 101 enzyme families fulfilled this criterion, and all homologous sequences from each expanded enzyme family were retrieved from the Genomes DB. The expanded enzyme families were classified into enzymes devoted to central metabolism, and expanded enzymes that may perform other functions, including NP biosynthesis. The latter classification was done using bidirectional best hits analyses, under the assumption that orthologous relationships imply the same function, and *vice versa*. However, at this stage, involvement of these enzymes in other metabolic functions, such as catabolism or resistance mechanisms, could not be ruled out.

We therefore, as our next step, aimed at identifying expanded enzymes that may have been recruited to catalyze conversions during NP biosynthesis. For this purpose, we performed a sequence similarity search between the Seed NP DB and the central metabolism DB obtained from the previous analyses. Recruitment events in NP BGCs were called when a sequence in an expanded enzyme family had a homologous sequence in the Seed NP DB. This search revealed that a total of 23 out of 101 expanded enzyme families had recognizable recruitment events by NP BGCs included in our Seed NP DB, which by no means represents an exhaustive catalog of NP pathways (Supplementary table 4). The remaining 75 expanded enzyme families may also include recruitment events by other NP BGCs not included in our Seed NP DB, or by other functional forms of metabolism such as degradation of xenobiotics. Nevertheless, the 23 enzyme families with recruitments analyzed in the following steps served as proof of concept for the discovery of new BGCs using phylogenomics.

To identify new recruitment events, we used phylogenetic analysis to define the evolutionary relationships in the 23 expanded-then-recruited enzyme families, and to infer possible functional fates for each homolog. In most of the 23 phylogenetic reconstructions (provided as online supplementary material), the central metabolic homologs formed clades with topologies that recapitulate speciation events. This could be confirmed by comparing the obtained topologies with a species phylogenetic tree constructed using neutral markers, such as the RNA Polymerase Beta subunit or RpoB (supplementary figure 1). In clear contrast, expansion events grouped in one or more independent clades, and the most divergent clades often included the recruited homologs from the Seed NP database (Figure 1B). We then assumed that high sequence divergence reflects rapid evolution, as expected for adaptive metabolism such as NP biosynthesis. Therefore, divergence of the clades that included homologs from the Seed NP DB was used to call positive hits (Figure 1B). Using these criteria we identified 515 enzyme hits contained in phylogenetic trees, which were visually inspected.

To further support the functional association of these enzyme’s hits with NP biosynthesis, we retrieved up to 80 Kbp of surrounding genomic sequence harboring the targeted genes (71.3 Kbp in average, 19.9 Kbp standard deviation). This process yielded 423 contigs out of the 515 enzyme hits, as some of these hits (almost 18%) were located in short contigs that could not be properly annotated. The 423 retrieved contigs were mined for BGCs of known classes of NPs using antiSMASH 3.0 (Weber et al, 2015), as well as for putative BGCs using ClusterFinder with a probability cut-off of 0.70 (Cimermancic et al, 2014). This analysis allowed us to “validate” our predictions after confirming that a hit enzyme was located within the boundaries of BGCs that could be predicted and annotated with these tools. For instance, 15% of the enzyme hits came from the Seed NP DB, whilst 59% coincided with AntiSMASH (43%), ClusterFinder (9%), or both (7%) (Figure 2A and supplementary table 5).

The remaining 26% of the enzyme hits were not included within the boundaries of BGCs predicted by neither AntiSMASH nor ClusterFinder, and therefore we refer to as “not validated”. Detailed analysis, however, revealed that 11% of the total hits, belonging to this category (not validated in Figure 2A), are at 20 Kbp or less away from predicted BGCs. This observation suggests that these enzymes escaped from the detection limits of the algorithms used, either due to their location at few Kbp beyond their boundary detection rules, or for having scores below the cut-off used for ClusterFinder predictions (0.70). The remaining 15% of the total hits, belonging to this category, could only be predicted by our approach (Figure 2A), and may represent truly novel BGCs. As a first step to look into this possibility, we analyzed the distribution and annotation of these hits, and we noticed that some enzyme families seem to have more predictive value as discussed further.

**Figure 2.**
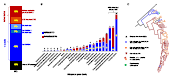
Overview of the BGC predictions obtained after the reconstruction of the evolution of NP biosynthesis. A. Distribution of validated and not validated enzyme hits. Most hits were validated by AntiSMASH and/or ClusterFinder. Refer to text for further details B. Distribution of hits per recruited enzyme family. The numbers show the total hits either validated or not validated for each enzyme family. C. Zoom-in of the sub-clade containing enzyme hits belonging to the 3-carboxyvinyl-phosphoshikimate transferase or AroA enzyme family. The complete tree, from which this sub-clade was extracted, can be seen in Figure 1B or Tree S4. Asterisks indicate enzymes recruited into known BGCs. blue branches and diamonds mark hits validated by other NP genome mining methods, orange triangles on orange branches mark not-validated hits located at less than 20 Kbp from a predicted BGC. Red X letters on red branches indicate not validated hits not related to predicted BGCs, red or orange circles indicate hits belonging to a family of new putative BGCs, finally green crosses indicates two experimentally characterized recruitment events discussed further in this article.

For instance, 43% of the total hits were found within three enzyme families, namely, asparagine synthase, 3-phosphoshikimate-1-carboxivinyl transferase and 2-dehydro-3-deoxyphosphoheptanoate aldolase (Figure 2B). This may be due to the limitations of our input databases. However, it is also tempting to speculate that enzyme recruitment is biased for certain enzyme families, at least more than currently acknowledged (Medema et al., 2015). Along these lines, the 3-phosphoshikimate-1-carboxivinyl transferase family, or AroA, comprise 38% of the enzyme hits that could not be validated, and includes the largest number of hits not associated to BGCs (Figure 2C). The not validated hits may represent the lowest confidence predictions, but also those with the highest potential for discovery of unprecedented chemistry (Medema and Fischbach, 2015).

The experimental characterization of this type of BGCs, i.e. not validated predictions, is challenging. Therefore, we focused on providing experimental evidence to validate our approach by selecting a predicted BGC as supported by other well-known NP enzymes, but which also includes an AroA hit that is unprecedented within the synthesis of NPs (Supplementary table 6). The selected prediction included known NP enzymes, namely, a two-gene PKS assembly, but also a 3-phosphoshikimate-1-carboxivinyl transferase homolog that is different to the AroA enzymes present in the BGCs of the polyketide asukamycin (Rui et al, 2010) and phenazines (Seeger et al, 2011), as discussed further.

### Discovery of arseno-organic metabolites related to a novel AroA enzyme

*S. coelicolor* A3(2) and *S. lividans* 66 are closely related model streptomycetes that have been thoroughly investigated both, before the genomics era and with genome mining approaches (Nett et al 2009, Bentley et al, 2002, Cruz-Morales et al, 2013). Presumably, most of their NPs biosynthetic repertoire has been elucidated (Challis 2014). Furthermore, several methods for genetic manipulation of these two strains are available (Kieser et al, 2000), making them ideal for proof of concept experiments. Specifically, our phylogenomics analysis led to five hits related to three BGCs in the *S. coelicolor* and *S. lividans* genomes (Supplementary table 5). Four of these hits (in both strains), included known recruitments that are associated with the BGC responsible for the synthesis of the calcium-dependent antibiotic or CDA (Hojati et al, 2002). The remaining hit, which caught our attention, is related to the AroA enzyme family, which catalyzes the transfer of a vinyl group from phosphoenolpyruvate (PEP) to 3-phosphoshikimate, forming 5-enolpyruvylshikimate-3-phosphate and releasing phosphate. The reaction is part of the shikimate pathway, a common pathway for the biosynthesis of aromatic amino acids and other metabolites (Zhang and Berti, 2006).

The phylogenetic reconstruction of the actinobacterial AroA enzyme family shows a major clade associated with central metabolism; this clade includes SLI_5501 from *S. lividans* and SCO5212 from *S. coelicolor*. More importantly, the phylogeny also includes a divergent clade with at least three sub-clades, two that include family members linked to the known BGCs of the polyketide asukamycin (Rui et al, 2010) and phenazines (Seeger et al, 2011); as well as AroA homologs from 26% of the genomes of our database. In *S. coelicolor* and *S. lividans*, these recruited homologs are encoded by SLI_1096 and SCO6819, and they are marked with green crosses in Figure 2C. These genes are situated only six genes upstream of the two-gene PKS system (SCO6826-7 and SLI_1088-9, respectively) used by AntiSMASH and ClusterFinder to “validate” these BGCs in the previous section (supplementary table 6). The PKS was identified in *S. coelicolor* since the early genome mining efforts in this organism, but often it was referred to as a cryptic BGC, which did not include these *aroA* homologs (Bentley et al, 2002; Nett et al, 2009).

Given that the aroA genomic context is highly conserved between the genomes of *S. lividans* and *S. coelicolor,* for simplicity we will refer to the *S. lividans* genes only (Supplementary table 6; MIBIG accession BGC0001283). The syntenic region spans from SLI_1077 to SLI_1103, including several biosynthetic enzymes, regulators and transporters, suggesting a functional association. Further annotation revealed the presence of a 2,3-bisphosphoglycerate-independent phosphoenolpyruvate mutase enzyme (SLI_1097; PPM), downstream and possibly transcriptionally coupled to the aroA homolog. Thus, a functional link between these genes, as well as with the phosphonopyruvate decarboxylase gene (PPD; SLI_1091) encoded in this BGC, was proposed. The combination of mutase-decarboxylase enzymes is a conserved biosynthetic feature of NPs containing Carbon-Phosphate bonds (Metcalf and van der Donk 2009), but that were not be detected by AntiSMASH or ClusterFinder (Supplementary table S6).

Other non-enzymatic functions within this BGC could be annotated, including a set of ABC transporters, originally annotated as phosphonate transporters (SLI_1100 and SLI_1101), and four arsenic tolerance-related genes (SLI_1077-1080) located upstream. These genes are paralogous to the main arsenic tolerance system encoded by the *ars* operon (Wang et al, 2006), which is located at the core of the *S. lividans* chromosome (SLI_3946-50). This BGC also codes for regulatory proteins, mainly arsenic responsive repressor proteins (SLI_1078, SLI_1092, SLI_1102 and SLI_1103). Thus, overall, our detailed annotation suggests a link between arsenic and phosphonate biosynthetic chemistry. Accordingly, in order to reconcile the presence of phosphonate-like biosynthetic, transporter and arsenic resistance genes, within a BGC, we postulated a biosynthetic logic analogous to that of phosphonate biosynthesis, but involving arsenate as the driving chemical moiety (Figure 3A).

Prior to functional characterization, the abovementioned hypothesis was further supported by the three following observations. First, arsenate and phosphate are similar in their chemical and thermodynamic properties, causing phosphate and arsenate-utilizing enzymes to have overlapping affinities and kinetic parameters (Tawfik and Viola, 2011; Elias et al, 2012). Second, previous studies have demonstrated that AroA is able to catalyze a reaction in the opposite direction to the biosynthesis of aromatic amino acids with poor efficiency, namely, the formation of PEP and 3-phosphoshikimate from enolpyruvyl shikimate 3-phosphate and phosphate (Zhang and Berti, 2006). Indeed, arsenate and enolpyruvyl shikimate 3-phosphate can react to produce arsenoenolpyruvate (AEP), a labile analog of PEP, which is spontaneously transformed into pyruvate and arsenate (Zhang and Berti, 2006). Third, it has been demonstrated that the phosphoenolpyruvate mutase, PPM, an enzyme responsible for the isomerization of PEP to produce phosphonopyruvate, is capable of recognizing AEP as a substrate. Although at low catalytic efficiency, the formation of 3-arsonopyruvate by this enzyme, a product analog of the phosphonopyruvate intermediate in phosphonate NPs biosynthesis (Metcalf and van der Donk, 2009), has been reported (Chawla et al, 1995).

Altogether, the previous evidence was used to postulate a novel biosynthetic pathway encoded by SLI_1077-SLI_1103, which may direct the synthesis of a novel arseno-organic metabolite (Figure 3A) (A detailed functional annotation, and biosynthetic proposal is provided in the supplementary table 6 and supplementary figure 2).

**Figure 3.**
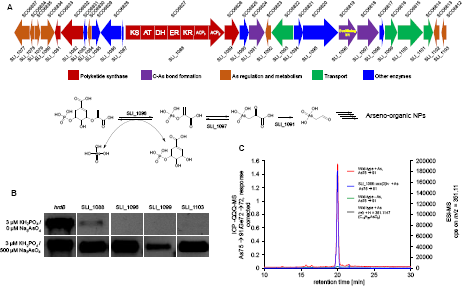
Discovery of a BGC for arseno-organic NPs in *S. coelicolor* and *S. lividans*. **A.** The conserved arseno-organic BGC in *S. lividans* 66 and *S. coelicolor* is shown. The proposed biosynthetic logic for early intermediates in the biosynthesis of arseno-organic metabolites is shown. **B.** Transcriptional analysis of selected genes within the arseno-organic BGC, showing that gene expression is repressed under standard conditions, but induced upon the presence of arsenate. **C.** HPLC-Orbitrap/QQQ-MS trace of organic extracts from mycelium of wild type and the SLI_1096 mutant showing the detection of arsenic-containing species. Three *m/z* signals were detected within the two peaks found in the trace from the wild type strain grown on the presence of arsenate. These *m/z* signals are absent from the wild type strain grown without arsenate, and from the SLI_1096 mutant strain grown on phosphate limitation and the presence of arsenate. Identical results were obtained for *S. coelicolor* and the SCO6819 mutant.

To determine the product of the predicted BGC, in both *S. lividans* and *S. coelicolor*, we used expression analysis, as well as comparative metabolic profiling of wild type and mutant strains. Using RT-PCR analysis, we first determined the transcriptional expression profiles of one of the PKS genes (SLI_1088), *aroA* (SLI_1096), one of the *arsR*-like regulator (SLI_1103), and the periplasmic-binding protein of the ABC-type transporter (SLI_1099). As expected for a cryptic BGC, the results of these experiments demonstrate that the proposed pathway is repressed under standard laboratory conditions. We then analyzed the potential role of arsenate as the BCG inducer upon the addition of a gradient concentration of arsenate (0 – 500 µM) and phosphate (3-300µM). Indeed, we found that the analyzed genes were induced when both *S. lividans* and *S. coelicolor* were grown in the presence of 500 µM of arsenic and 3 µM of phosphate (Figure 3B).

In parallel, we used PCR-targeted gene replacement to produce mutants of the SLI_1096 and SCO6819 genes, and analyzed the phenotypes of the mutant and wild type strains on a combined arsenate/phosphate gradient, i.e. low phosphate and high arsenate, and *vice versa*. Intracellular organic extracts were analyzed using HPLC coupled with an ICP-MS calibrated to detect arsenic-containing molecular species. Simultaneously, a high-resolution mass spectrometer determined the mass over charge (*m*/*z)* of the ions detected by the ICP. This set up allows for high-resolution detection of arseno-organic metabolites (Amayo et al, 2011). Using this approach, we detected the presence of an arseno-organic metabolite in the organic extracts of both wild-type *S. coelicolor* and *S. lividans*, with *m*/*z* value of 351.1147. These metabolites could not be detected in either identical extracts from wild type strains grown in the absence of arsenate or in the mutant strains deficient for the SLI_1096/SCO6819 genes (Figure 3C). Thus, the product of this pathway may be a relatively polar arseno-polyketide. The actual structures of these products are still subject to further chemical investigation and will be discussed in detail in a future publication.

### Closure of the conceptual loop: Genome mining for arseno-organic BGCs

Confirmation of a link between SLI_1096/SCO6819 and the synthesis of an arseno-organic metabolite provides an example on how genome-mining efforts, based in novel enzyme sequences, can be advanced. For instance, co-occurrence of divergent SLI_1096 orthologs, now called arsenoenolpyruvate synthases (AEPS); SLI_1097 arsenopyruvate mutase (APM) and arsonopyruvate decarboxylase (APD), can be now confidently used beacons to mine bacterial genomes. Indeed, 26 BGCs with the potential to synthesize arseno-organic metabolites, all of them encoded in genomes of myceliated *Actinobacteria*, were identified after sequence similarity searches using the non-redundant GenBank database (Figure 4). The divergence and potential chemical diversity within these arseno-related BGCs was analyzed using phylogenetic reconstructions with a matrix of three conserved genes of these BGCs. This analysis suggests three possible sub-classes with distinctive features, PKS-NRPS independent, PKS-NPRS-hybrid dependent and PKS-dependent biosynthetic systems.

**Figure 4.**
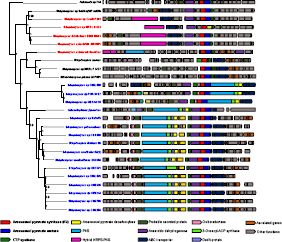
Novel BGCs for arseno-organic metabolites found in *Actinobacteria*. The BGCs were found by mining for the co-occurrence of arsenoenol pyruvate synthase (AEPS), arsenopyruvate mutase (APM). Related BGCs were only found in actinomycetes. The phylogeny was constructed with a concatenated matrix of conserved enzymes among the BGCs including AEPS, APM and CTP synthase. Variations in the functional content of the BGCs are accounted by genes indicated with different colors. Three main classes or arseno-organic BGCs could be expected from this analysis: PKS-NRPS independent (black names), PKS-NPRS-hybrid dependent (red names) and PKS-dependent biosynthetic systems (blue names).

In addition, once a hit predicted to perform an unprecedented function is positively linked to a new BGC and its resulting metabolite, this could lead to identification of novel classes of conserved BGCs (Figure 2C). To illustrate this, the non-validated AroA hits within the NP-related clade of this phylogenetic tree were manually curated in search for a conserved BGC. One particular case, representing a previously unnoticed sub-clade that belongs to the not validated category was found to appear frequently. After detailed annotation, it was found that this BGC is conserved in at least sixty-three *Streptomyces* genomes, and in the genome of *Microtetraspora glauca* NRRL B-3735, which are included in the non-redundant GenBank database (supplementary table 7). For these analyzes, 15 genes upstream and downstream the AroA hit were extracted as before and annotated on the basis of the locus from *Streptomyces griseolus* NRRL B-2925 (Supplementary table 8 and supplementary figure 3). Indeed, this locus has some of the expected features for an NP BGC, including: (i) gene organization suggesting an assembly of operons, most of them transcribed in the same direction; (ii) genes encoding for enzymes, regulators and potential resistance mechanisms; and (iii) enzymes that have been found in other known NP BGCs. The latter observation, which actually includes seven genes out of 31, present in an equal number of NP BGCs, actually supports the NP nature of this locus.

Overall, we conclude that functional predictions based upon phylogenomics analysis has great potential to ease natural product and drug discovery by means of accelerating the conceptual genome-mining loop that goes from novel enzymatic conversions, their sequences, and propagation after homology searches. The proposed approach now makes the retrieval of arseno-organic BGCs possible that were previously not detected by AntisMASH or ClusterFinder (for instance the one from *Streptomyces ochraceiscleroticus*). The current work provides the proof of concept and the fundamental insights to develop a bioinformatics pipeline based upon evolutionary principles, to be referred as EvoMining.

## Acknowledgements

We are indebted with Marnix Medema, Paul Straight and Sean Rovito, for useful discussions and critical reading of the manuscript, as well as with Alicia Chagolla and Yolanda Rodriguez of the MS Service of Unidad Irapuato, Cinvestav, and Araceli Fernandez for technical support in high-performance computing. This work was funded by Conacyt Mexico (grants No. 179290 and 177568) and FINNOVA Mexico (grant No. 214716) to FBG. PCM was funded by Conacyt scholarship (No. 28830) and a Cinvestav posdoctoral fellowship. JF and JFK acknowledge funding from the College of Physical Sciences, University of Aberdeen, UK.

## Competing interests

PCM and FBG have filed patent applications related to this work. The other authors declare that no competing interests exist.

